# Molecular architecture of the TasA biofilm scaffold in *Bacillus subtilis*

**DOI:** 10.1101/2022.03.14.484220

**Authors:** Jan Böhning, Mnar Ghrayeb, Conrado Pedebos, Daniel K. Abbas, Syma Khalid, Liraz Chai, Tanmay A. M. Bharat

## Abstract

Many bacteria in nature exist in multicellular communities termed biofilms. Cells within biofilms are embedded in a primarily self-secreted extracellular polymeric matrix that provides rigidity to the biofilm and protects cells from chemical and mechanical stresses. In the Gram-positive model biofilm-forming bacterium *Bacillus subtilis*, TasA is the major protein component of the biofilm matrix, where it has been reported to form functional amyloid fibres contributing to biofilm structure and stability. The structure of TasA fibres, however, and how fibres scaffold the biofilm at the molecular level, is not known. Here, we present electron cryomicroscopy structures of TasA fibres, which show that rather than forming amyloid fibrils, TasA monomers assemble into filaments through donor strand complementation, with each subunit donating a β-strand to complete the fold of the next subunit along the filament. Combining electron cryotomography, atomic force microscopy, and mutational studies, we show how TasA filaments congregate in three dimensions to form abundant fibre bundles that are essential for *B. subtilis* biofilm formation. This study explains the previously observed biochemical properties of TasA and shows, for the first time, how a bacterial extracellular globular protein can assemble from monomers into β-sheet-rich fibres, and how such fibres assemble into bundles in biofilms. We establish a hierarchical, atomic-level assembly mechanism of biofilm scaffolding that provides a structural framework for understanding bacterial biofilm formation.

## Introduction

Biofilms are the primary mode of microbial multicellular existence in nature [1]. Biofilms form on natural surfaces in the environment as well as inside the bodies of living organisms during infection processes, and are further commonly found on abiotic surfaces such as catheters and medical devices [2]. In biofilms, bacterial cells encase themselves in a matrix of extracellular polymeric substance (EPS), made primarily of filamentous polymeric molecules, including proteins, polysaccharides, and DNA [3]. The EPS matrix protects cells against physical and chemical stresses including antibiotic treatment [1-5] and is a hallmark of all bacterial biofilms. Therefore, understanding how polymeric molecules assemble in the extracellular matrix (ECM), and how they serve to scaffold biofilms, is of fundamental importance. Nevertheless, to the best of our knowledge, no high-resolution structures of biofilm scaffold fibres are available to help bridge this mechanistic gap in our understanding of biofilm formation.

The soil bacterium *Bacillus subtilis* is one of the best-studied model organisms for investigating biofilm formation [6-12]. The *B. subtilis* biofilm ECM is rich in exopolysaccharide and protein components, the major proteinaceous component being a 26 kDa fibre-forming protein called TasA [13, 14]. TasA is expressed as part of the *tapA*-*sipW-tasA* operon, which is controlled by the biofilm regulator protein SinR [15, 16]. TasA was originally characterised as a spore coat-associated protein with antimicrobial activity, deletion of which results in defects in spore coat structure [14]. Later studies revealed that TasA assembles into fibres that form the major component of the *Bacillus subtilis* biofilm matrix [13], with deletions of *tasA* resulting in significant defects in biofilm and pellicle formation that can be rescued by addition of exogenous TasA [17]. The *tasA* operon further encodes accessory proteins; TapA, a minor matrix component often found associated with TasA, which is thought to be involved in anchoring TasA to the cell wall and in accelerating its assembly [18, 19], and the membrane-associated signal peptidase SipW that cleaves off the signal peptides of TasA and TapA [20].

An X-ray atomic structure of monomeric TasA has been solved showing a globular subunit, with a classical Jellyroll fold [21]. Interestingly, TasA fibres have been shown to possess several properties characteristic of amyloids, including high β-sheet content, resistance to depolymerisation and mechanical disruption, binding to the amyloid-specific A11 antibody, and staining positive in Congo Red and Thioflavin T assays [13, 17, 21-23], suggesting that TasA belongs to a growing list of so-called functional amyloid fibres [24], which also include biofilm matrix proteins in several Gram-negative bacteria such as *Escherichia coli* [25] and *Pseudomonas fluorescens* [26]. In contrast, a recent study proposed that TasA fibres consist of a linear arrangement of globular subunits that does not involve structural rearrangements in the transition of TasA monomers to fibres, suggesting that fibres formed by recombinant TasA are not amyloid [27]. Moreover, a recent X-ray diffraction study showed that TasA fibrils only possess a weak cross-β-sheet pattern [11]. Given a lack of direct structural data, it is not possible to relate these previous observations to the underlying molecular arrangement of TasA.

In this study, we address these ambiguities and bridge the gap in our mechanistic understanding of how abundant biofilm matrix proteins scaffold biofilms using electron cryomicroscopy (cryo-EM) structure determination, electron cryotomography (cryo-ET) and atomic force microscopy (AFM) of purified single fibres, reconstituted fibre bundles, and biofilms respectively. Combining our structural data with molecular dynamics (MD) simulations and mutational studies, we provide a structural basis of biofilm scaffolding by the TasA protein in the Gram-positive model organism *B. subtilis*.

## Results

### Atomic structure of TasA fibres

To gain insights into the structure of TasA fibres, we purified TasA from *B. subtilis* using previously published procedures [28]. Following cryo-EM sample preparation, we observed highly ordered fibres with a characteristic periodic appearance on cryo-EM grids (Figure 1A), consistent with previous EM studies [27, 29]. We performed helical reconstruction from cryo-EM images and determined a 3.5 Å resolution structure of TasA fibres (Figures 1B-1D; Figures S1 and S2). To our surprise, this structure shows that TasA monomers are held within filaments by a key inter-subunit interaction provided by an N-terminal β-strand that extends from each subunit into the next, where it completes the β-sheet-rich fold of TasA in each subunit (Figures 1B-1D), thus providing an extensive inter-subunit interaction. In addition to the β-sheet hydrogen bonding of the donor strand with the next subunit, there are extensive hydrophobic interactions between donor strand residues (F29, I32, F39) and the next two subunits, i.e. the donor strand of the (n-1)_th_ subunit interacts with the n_th_ and the (n+1)_th_ subunits (Figures 1D and S2). The C-terminal residues 239-261, previously found to be unstructured in NMR studies of monomeric TasA and unresolved in the monomer X-ray structure [21], are ordered in the fibre structure and form another inter-subunit interface (Figure S2B). Thus, the filament structure is neither representative of cross-β amyloids, nor is it a head-to-tail arrangement of globular subunits as previously suggested, but shares features of both, including inter-subunit interactions mediated by β-strands and an intact core fold of the monomer. The substantial extent of interactions between TasA subunits is likely responsible for the significant stability of the fibres previously observed [17], and the β-sheet-rich architecture and extensive β-strand interactions between subunits may offer an explanation for why, even though TasA filaments do not form an amyloid structure [30], they show several characteristics of amyloids [17, 19].

**Figure 1.**
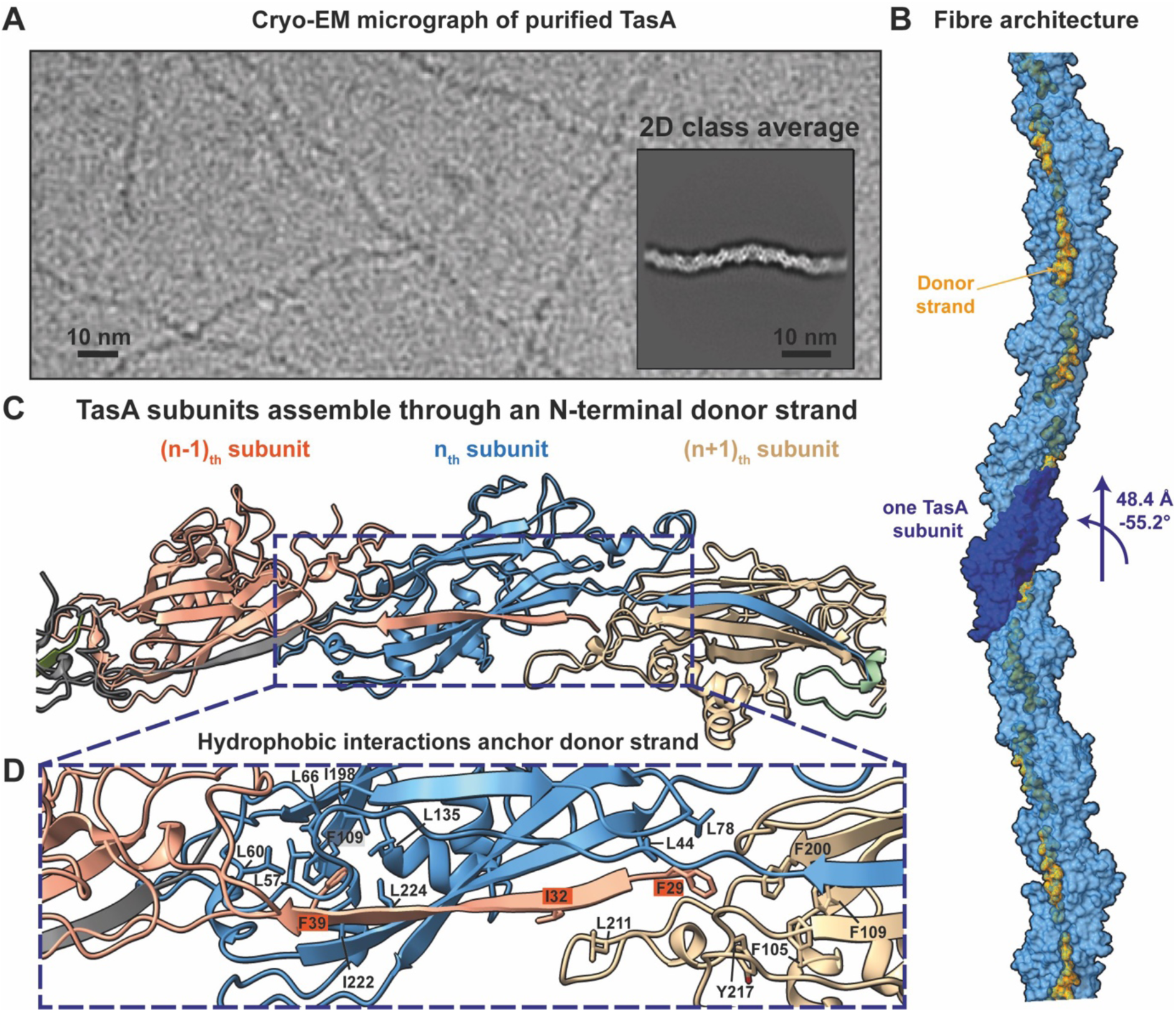
TasA polymerises through donor strand complementation. **(A)** Cryo-EM image of purified TasA shows fibres with a characteristic periodic appearance. The inset shows a two-dimensional class average of TasA fibres. **(B)** Arrangement of TasA subunits in fibres. One subunit is highlighted in a darker shade of blue, with the N-terminal donor strands in orange. **(C)** TasA subunits polymerise through an N-terminal donor strand completing the β-sandwich fold of the following (n+1)_th_ subunit. **(D)** Hydrophobic residues in the (n-1)_th_ subunit mediate donor strand interactions with the (n)_th_ and (n+1)_th_ subunits.

Remarkably, the subunit interactions through a donor β-strand observed in TasA fibres are reminiscent of Gram-negative bacterial pilus assembly systems (Type I and V pili, Figure S3A) that are based on donor-strand exchange [31, 32]. Interestingly, a β-sandwich fold is not only observed in such pilins, but also in non-bacterial proteins undergoing β-strand exchange, such as the Mcg IgG protein involved in light-chain amyloidosis (Figure S3B) [33], which together suggest that a β-sandwich fold may be especially amenable for facilitating donor strand complementation for the formation of fibrous assemblies.

### Rearrangement of N- and C-termini drives TasA fibre formation

The donor β-strand shields a significant hydrophobic groove in the fold of TasA (Figure S2D) that would otherwise be exposed if not complemented by a donor strand. It is, however, known that the monomeric form of TasA protein is soluble and stable [21, 28], indicating that residues are differently arranged between the monomeric and filamentous form of TasA. In order to analyse the structural differences in these two forms of TasA, we compared the monomeric, soluble form of TasA (PDB 5OF2) with the fibre form solved in this study (Figures 2A-2B). Remarkably, we observed that the binding site of the donor strand is self-complemented by a different β-strand (residues 48-61) in the monomeric form (Figures 2A-2B). In the fibre form, the previously self-complementing β-strand in monomeric form is displaced by the donor strand of the (n-1)_th_ subunit, and folds over the opposite β-sheet. Correspondingly, residues 39-47 are extended towards the next (n+1)_th_ subunit in the fibre form, presenting the donor strand (residues 28-38), allowing the next subunit to assemble (Figure 2A). The C-terminus of TasA is well-resolved in the fibre structure (Figures 2A and 2C; Figure S2B), suggesting a role of the C-terminal residues in promoting fibre stability.

**Figure 2.**
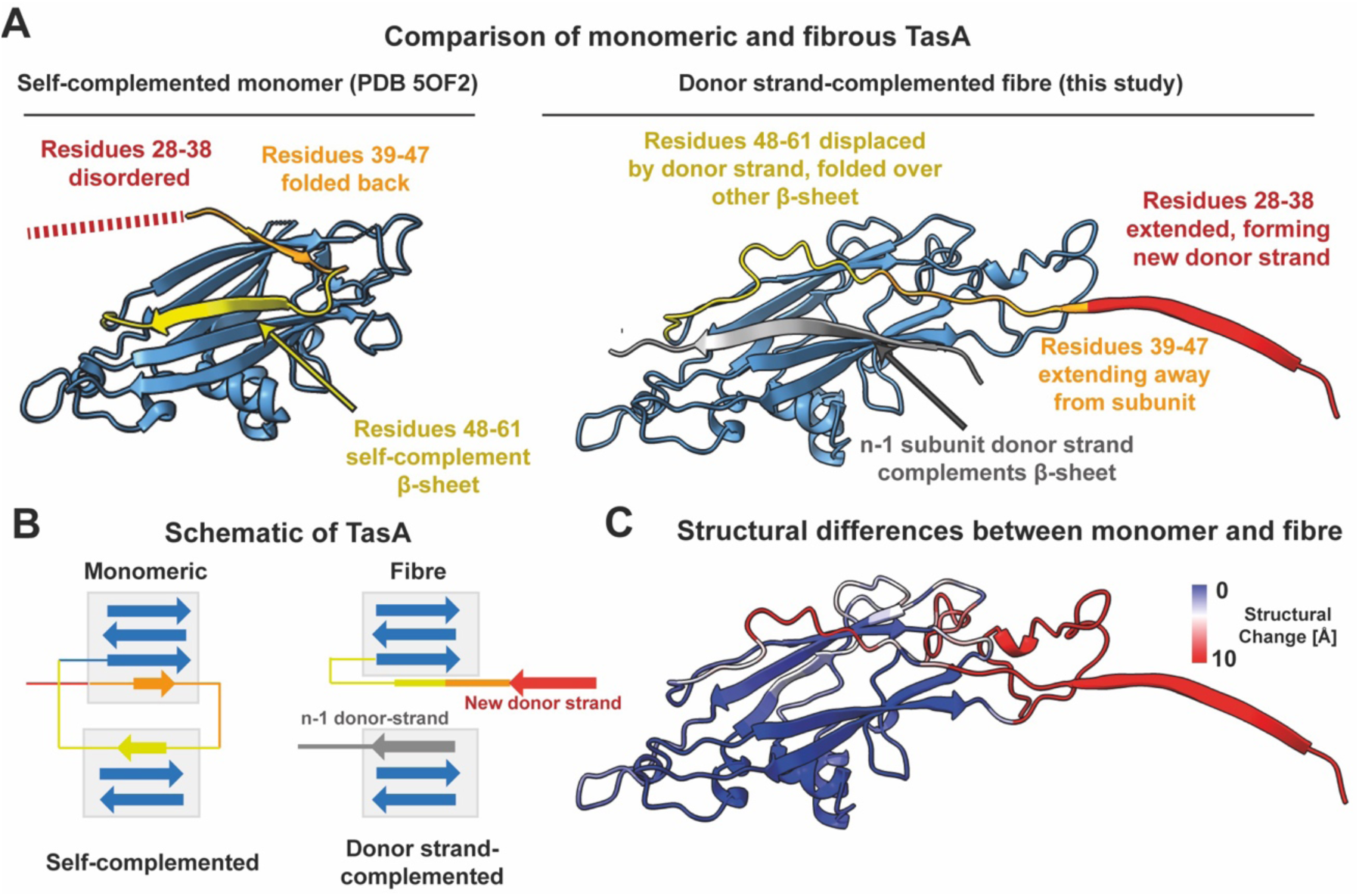
TasA monomers undergo re-arrangement at the N- and C-termini upon fibre formation. **(A)** Structural comparison of soluble, monomeric TasA (PDB 5OF2, left) and fibrous TasA (right). Segments of the N-terminus are coloured in yellow (residues 48-61), orange (residues 39-47) and red (residues 28-38) to facilitate visualisation. The C-terminus (residues 239-261) in the monomeric, self-complemented TasA (PDB 5OF2) is unstructured and thus not shown. **(B)** Schematic of β-sheet architecture (not to scale) demonstrating the structural differences between the monomeric and fibrous TasA. Same colours as in panel A are used. **(C)** Cryo-EM structure of TasA (fibre-form) coloured based on structural deviations from the monomeric form. Large deviations are observed at the N- and C-termini. Regions unstructured in monomeric TasA are coloured red.

Our structure suggests that, unless a TasA subunit is donor strand-complemented, it cannot extend its own donor strand as this would otherwise expose a hydrophobic groove (Figure S2D). Previous studies have shown that the accessory protein TapA nucleates TasA fibre formation in native cellular systems [19], and that it accelerates the assembly of TasA into fibres *in vitro* [18]. We noticed significant sequence similarity between the TapA N-terminal residues and the TasA donor strand (Figure S4), suggesting TapA might extend a donor strand itself. Since small, sub-stoichiometric subunits of protein fibre tips near the cell surface are not amenable to helical cryo-EM reconstruction, we employed AlphaFold-Multimer [34] to predict whether TapA could complement a TasA subunit. Indeed, the structural model shows the N-terminal TapA residues extending into the β-sandwich domain of TasA, completing the fold (Figure S4). Together with previous results showing that TapA nucleates fibre formation [18, 19], this suggests that TapA may promote the donor strand exchange reaction by providing the first donor strand during fibre formation in biofilms. This also provides a structural explanation for a previous observation that only a fraction of N-terminal residues of TapA are required for biofilm structuring by TasA [35].

### Interactions between TasA fibres

To obtain a structural understanding of how TasA fibres form the bundles observed in biofilms [22], we performed cryo-ET analysis on TasA bundles reconstituted *in vitro* (Figure 3). While we observed TasA fibres associated into large bundles, smaller bundles and even single fibres could consistently be detected, implying a low-affinity interaction between fibres. This observation is consistent with previous studies showing that TasA bundling is concentration-dependent [29], possibly relying on avidity effects for assembly. TasA filaments within the bundle were not randomly arranged, but rather orientationally aligned (Figure 3), which is a classical instance of phase separation into a nematic phase, also known as tactoid formation.

**Figure 3.**
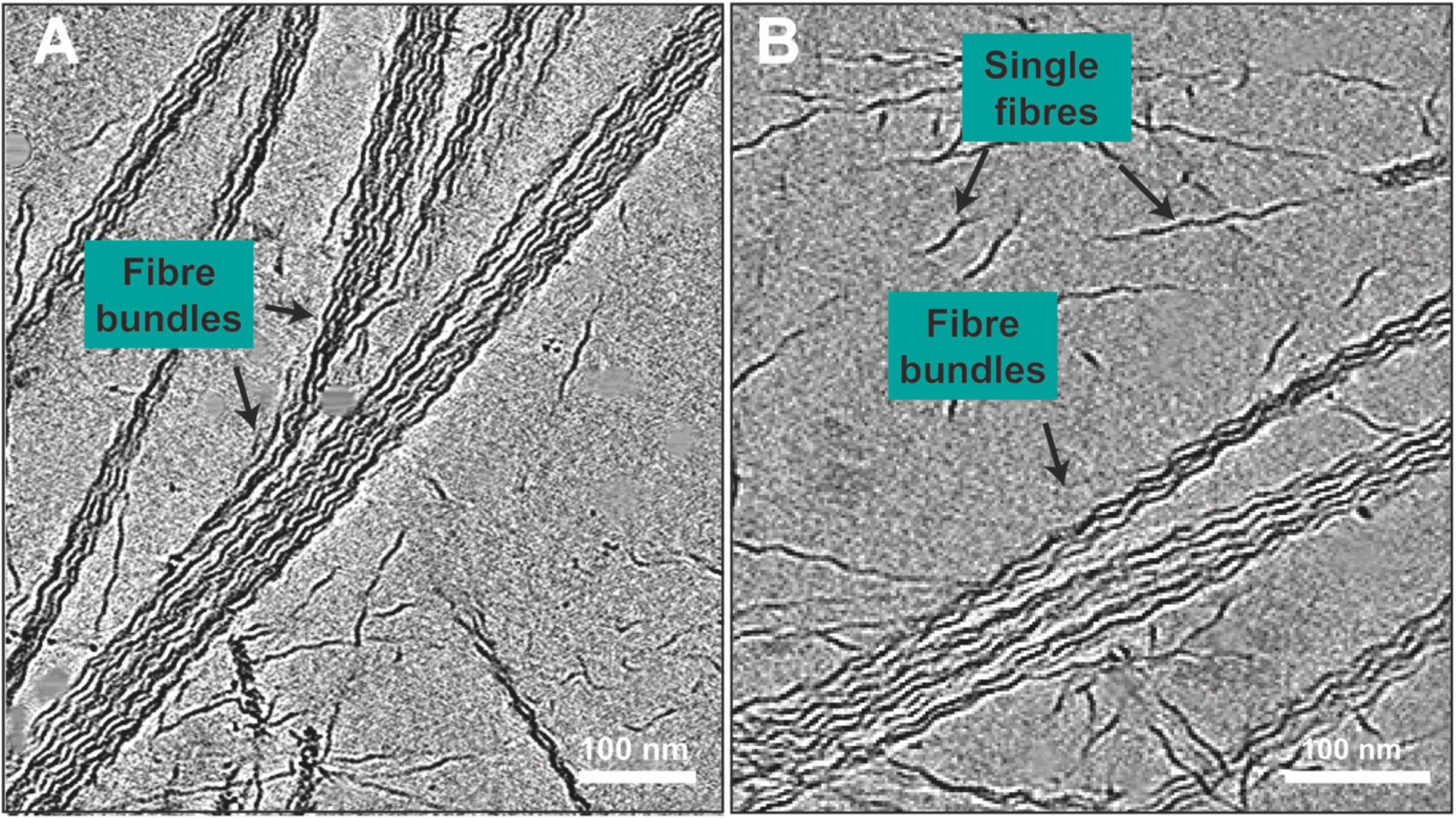
TasA fibres spontaneously associate into phase-separated bundles. **(A-B)** Cryo-ET slices of purified TasA shows the formation of locally ordered filament bundles. Both bundles and single fibres can be seen in the specimen (marked).

To probe the mode of interaction between fibres, we utilized cryo-ET data from doublets of TasA fibres, which represent the simplest interaction between fibres (Figure 4A-4C), and which are technically amenable for cryo-ET-based modelling [36]. The atomic model of a TasA fibre was docked into the cryo-ET density (Figures 4B and 4C). Our atomised model of the TasA doublet shows that fibres do not interact consistently all along the length of the doublet, but rather periodically every ∼32 nm. Our fits into multiple tomograms consistently place a loop extending from the fibre core (residues 174-177 with sequence DGKT, Figure S5) at the fibre-fibre interface (Figure 4D), which suggests a role of these loop residues in promoting bundle formation. Simple geometrical considerations support this hypothesis, as the loop is the structural element that extends furthest out from the TasA fibre, ideally placed for mediating such an interaction (Figure 4E).

**Figure 4.**
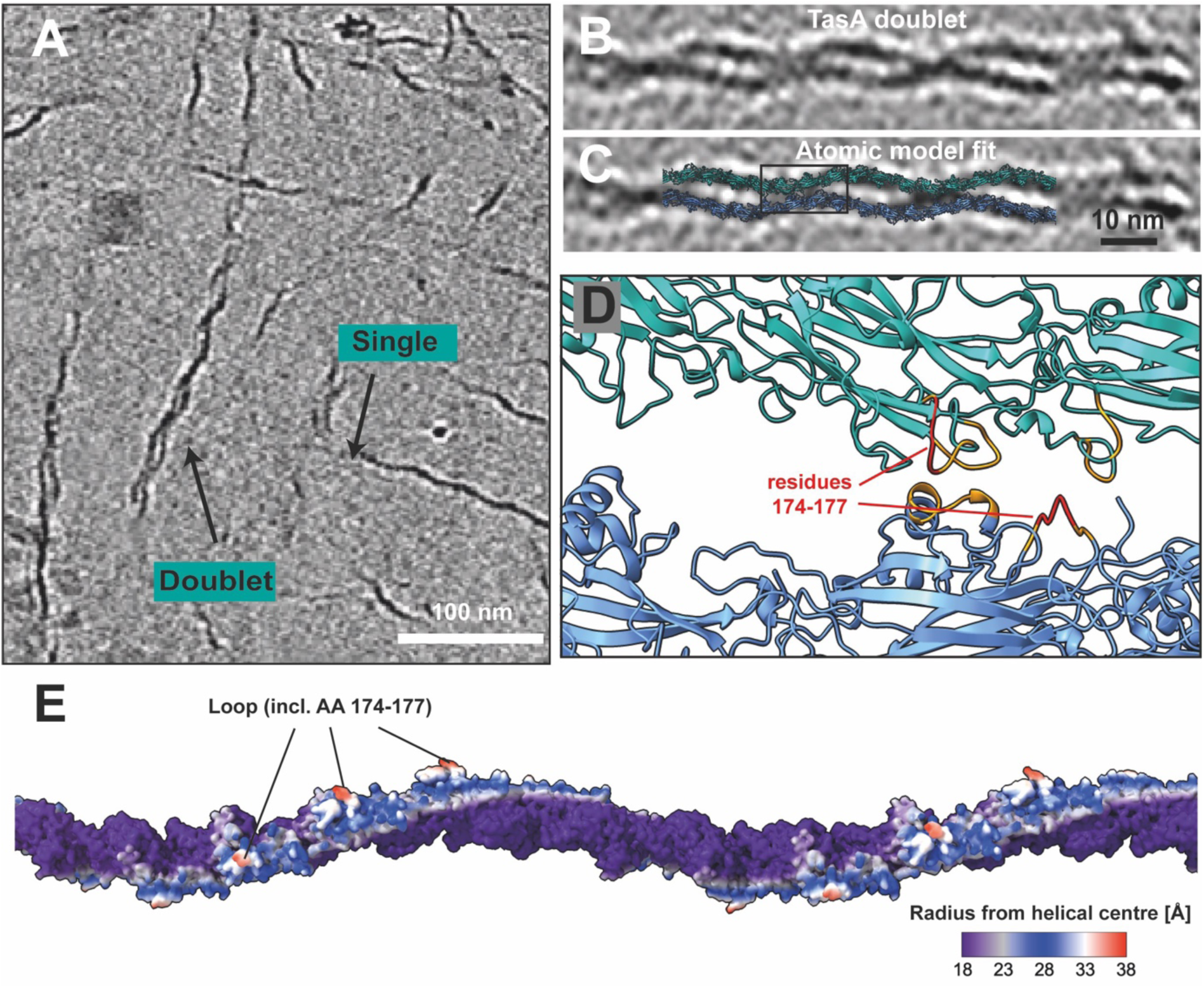
TasA fibres form bundles through periodic interactions with neighbouring fibres mediated by distal loop residues. **(A)** Tomographic slice showing TasA bundles, with the smallest bundles formed by two TasA fibres marked. **(B)** Cryo-ET data depicting the interaction of two TasA fibres, revealing that fibre-fibre interactions are periodic. **(C)** Fit of atomic models of two TasA fibres into the cryo-ET density. **(D)** Zoomed view of the atomised model verified by MD simulations. Residues found to be interacting are marked (orange), with the key residues 174-177 marked in red. (E) Atomic model of a TasA fibre (18 subunits shown) coloured by radius from the helical axis. The part of the filament extending out farthest (red) corresponds to the loop containing residues 174-177.

To determine whether fibre-fibre interactions are mediated through this part of the protein, we performed molecular dynamics (MD) simulations on a two-fibre system (Figure S5). The simulations confirmed a stable interaction that includes both the extended loop region (residues 173-179, IDKGTAP) as well as some additional residues at the fibre-fibre interface (Figure S5), including in a short α-helical segment (residues 153-160, IKKQIDPK) (Figure 4D). To further verify our structural observations, we generated genomic *B. subtilis* mutants expressing TasA in which residues 174-177 are mutated to poly-alanine (174-177_AAAA_), which, according to our model, would disrupt fibre-fibre bundling. Biofilms and pellicles with this mutation (174-177_AAAA_) showed an aberrant morphology compared to wild-type *B. subtilis* biofilms, reminiscent of a Δ*tasA* strain (Figures 5A-5C; Figure S6), strongly indicating that the bundling ability of TasA had been disrupted, supporting the inferences from our cryo-ET and MD based doublet model. Also, Cryo-EM of the 174-177_AAAA_ mutant fibres showed markedly reduced bundling (Figures S6D-S6E).

**Figure 5.**
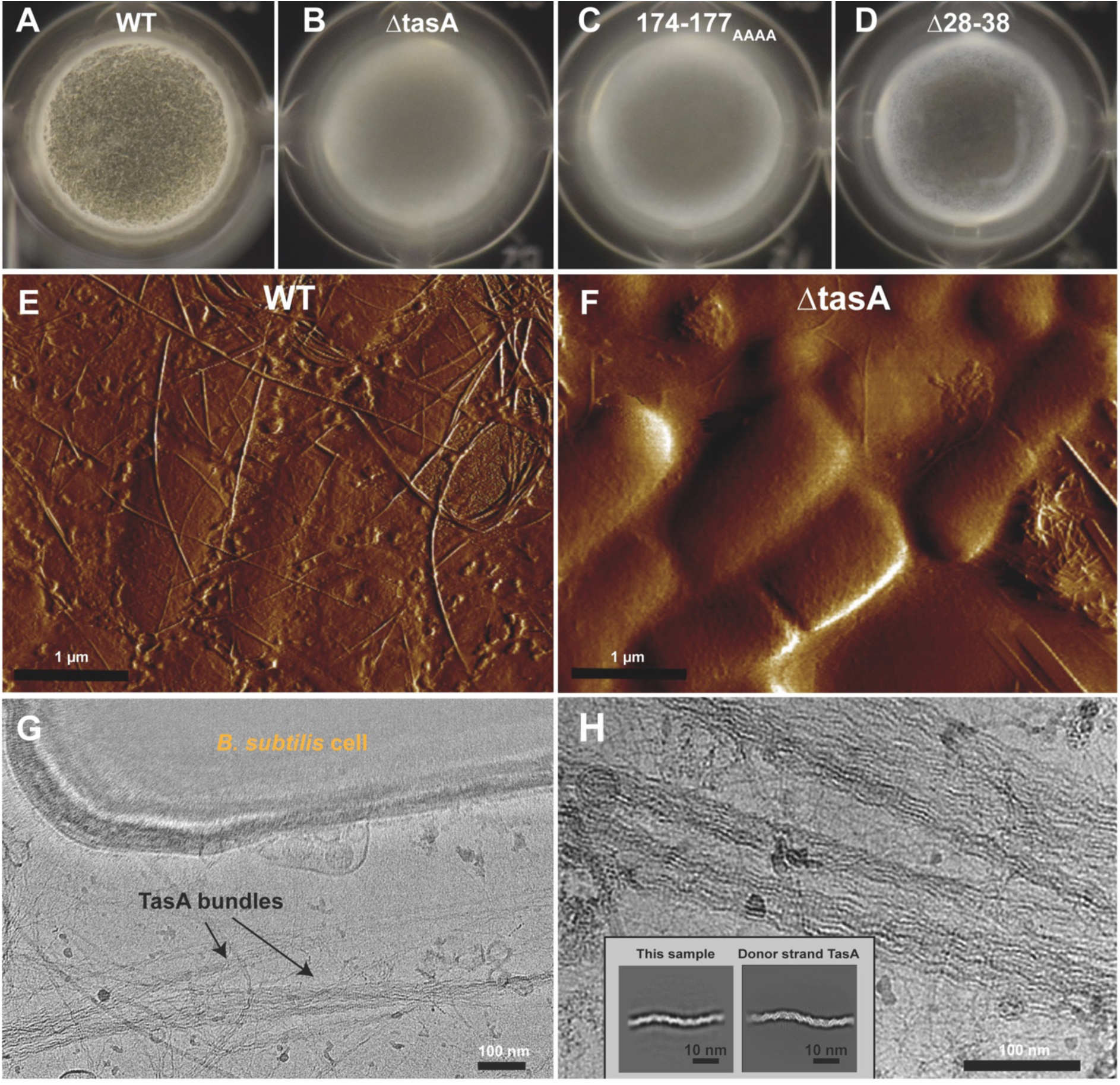
TasA in *B. subtilis* biofilms. **(A-D)** *B. subtilis* pellicles after a 24 hour incubation of strains: (A) wild-type, (B) Δ*tasA*, (C) fibre-fibre interface mutant (174-177_AAAA_) *tasA* and (D) donor strand mutant (Δ28-38) *tasA*. **(E-F)** AFM imaging comparing (E) wild-type (ZK5041) and (F) Δ*tasA* (ZK3657) *B. subtilis* pellicles shows an extensive interconnected network of TasA fibres stabilising the ECM. **(G)** Cryo-EM image of the ECM components of a *B. subtilis* Δ*eps* Δ*sinR* biofilm show bacterial cells along with TasA fibres and bundles. **(H)** TasA fibres in the ECM show the same structural features as purified TasA. Inset: Two-dimensional averaging shows that ECM biofilm and purified TasA are indistinguishable in their helical arrangements.

Additionally, we generated mutant strains expressing donor strand-truncated *tasA* (Δ28-38), which were found to form significantly defective biofilms and pellicles, showing an aberrant morphology compared to wild-type *B. subtilis* (Figures 5D and S6A). Cryo-EM of Δ28-38 TasA fibres showed defective fibre formation in TasA isolates (Figures S6D and S6F), consistent with the role of the donor strand in fibre assembly suggested by our structural data. These observed effects underline that both fibre formation and bundle formation of TasA are integral for *B. subtilis* biofilms and that their absence could not be compensated by other matrix components.

Next, by imaging intact wild-type pellicle biofilms using atomic force microscopy (AFM), we observed TasA forming networks of filaments adhered to cells that are not present in Δ*tasA* biofilms (Figures 5E-5F; Figure S7). This further confirms the role of TasA in scaffolding biofilms by forming an integral part of the ECM. We subsequently went on to test whether the fibres observed in biofilms correspond to those formed from purified TasA to rule out artefacts of *in vitro* preparation or a possible polymorphism of TasA fibres. Accordingly, we performed cryo-EM imaging of the ECM components of *B. subtilis* biofilms to examine the morphology of native TasA fibres. For this experiment, we used biofilms of an exopolysaccharide-deficient *B. subtilis* mutant (Δ*sinR* Δ*eps*), which, compared to wild-type *B. subtilis*, readily disassembled into its components upon gentle agitation and deposition on cryo-EM grids (Figures 5G-5H). A visual inspection of the specimen showed deposited *B. subtilis* cells together with components of the ECM (Figure S8). We observed that a major component of the specimen consisted of TasA fibres, occurring predominantly in fibre bundles (Figures 5G-5H; Figure S8), in line with previous reports [22]. Both fibre bundles and individual TasA fibres were observed on the grid, and strikingly, they were found to be identical to purified TasA (Figure 5H, inset), including the same subunit arrangement and helical repeat, strongly suggesting that the donor strand-exchanged TasA is the primary form of the protein in biofilms. It is worth noting that, compared to wild-type biofilms, the Δ*eps* biofilms disassemble easily into cells and matrix components, suggesting a complementary role of the matrix polysaccharides in promoting cohesion of the ECM.

## Discussion

The atomic structure of TasA fibres shows how this protein retains characteristics of the monomeric protein [21], which would be expected in a linear arrangement of globular subunits [27]. At the same time, β-strand interactions between TasA subunits in the filament result in a highly inter-connected and stabilized structure, as found in classical amyloids [13, 17, 21-23]. These unique features explain why previous studies have arrived at different expected subunit arrangements of TasA within the filament. While it is well established that functional amyloids are abundant in bacteria and beyond [24], we propose that other suggested functional amyloids could also exhibit similar characteristics, and that there may be a continuum of fibre-forming proteins - on one end arranged as classical amyloids [25, 37], to amyloid-like fibres displaying features shared with filaments formed from globular proteins on the other end [27].

The structure of TasA fibres is the first described case of a donor strand exchange system in Gram-positive bacteria. While Gram-negative systems [31, 32] do not appear to be evolutionarily related to the TasA/TapA system, the donor strand exchange suggests a common structural solution to the challenge of forming secreted protein fibres with high stability in the harsh and relatively uncontrolled extracellular space. We believe that this is a striking exemplar of convergent evolution, since the mechanisms of fibre assembly of TasA and Gram-negative pili are dramatically different. Compared to the well-characterised Gram-negative bacterial systems such as Chaperone-Usher Pathway (Type I) pili and Type V pili, TasA fibre assembly does not involve chaperones, usher proteins or proteolytic cleavage to promote transition into donor strand-complemented fibres. Further studies into the TasA/TapA system will be needed to understand in detail the mechanisms of assembly of TasA fibres as compared to Gram-negative pili.

In this study, we describe the first structure of a biofilm matrix fibre scaffold, important for understanding the arrangement of the ECM of bacterial biofilms. In particular, we show at the individual amino acid level how TasA monomers and fibres assemble and stack onto each other to form three-dimensional bundles, scaffolding *B. subtilis* biofilms (Figure 6). Our mutational, cryo-EM and AFM experiments together confirm that donor strand-exchanged TasA fibres form a large and crucial part of the *B. subtilis* biofilm ECM, and also confirm that TasA’s role as a scaffold is vital for the formation of rigid biofilms. This study thus establishes the structural basis of how filamentous fibres form a scaffold in the ECM of bacterial biofilms, with potential implications on other such systems including fibrous proteins that form bundles including Curli from *E. coli* [25], Fap from Pseudomonads [26] and phenol-soluble modulins and Bap from *Staphylococcus aureus* [38, 39].

**Figure 6.**
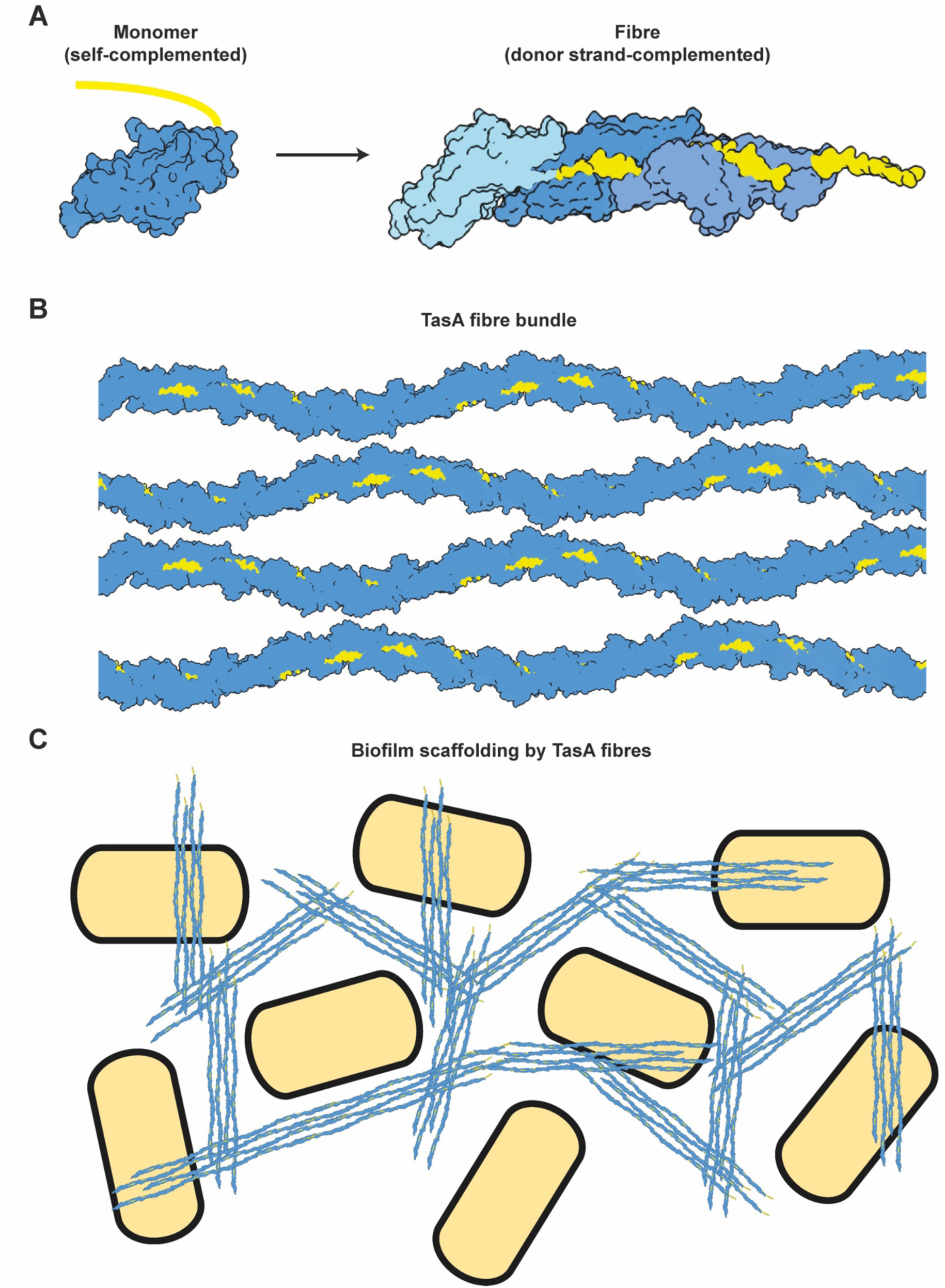
Schematic model of biofilm scaffolding by TasA. **(A)** Formation of fibres by transitioning from a self-complemented monomer to a donor-strand complemented fibre. **(B)** Bundle formation through periodic interactions of TasA fibres. **(C)** Biofilm scaffolding through formation of TasA networks.

A recurring theme in bacterial biofilms is the abundance of filamentous molecules including polysaccharides [40, 41], proteins [24] and DNA [42] in the ECM [3], which are possibly important for the formation of a phase-separated environment protecting cells within biofilms [43, 44]. Further studies into such mechanisms will be important to understand how the ECM is formed, which is of fundamental importance in comprehending how biofilms are built. Since the absence of the *B. subtilis* eps facilitated disassembly of the biofilm (Figure 5G-5H), this study provides another example of a synergy between filamentous proteins and ECM polysaccharides, described in other model bacterial systems including pathogenic species such as *Vibrio cholerae* and *Pseudomonas aeruginosa* [40, 41]. Taken together with previous studies, our data suggest that ordered, local interactions between filamentous molecules leading to phase-separation in the ECM might be a general organisational principle for biofilm formation.

## Acknowledgments

The authors would like to thank Sjors Scheres and Jan Löwe for helpful discussions, and are grateful to Sigal Ben-Yehuda and to Bing Zhou for their generous help with strain construction. T.A.M.B. is a recipient of a Sir Henry Dale Fellowship, jointly funded by the Wellcome Trust and the Royal Society (202231/Z/16/Z). T.A.M.B. would like to thank the Vallee Research Foundation, the Leverhulme Trust and the Lister Institute of Preventative Medicine for support. J.B. is supported by a Medical Research Council graduate studentship (grant numbers MR/K501256/1 and MR/N013468/1). M.G. acknowledges the support of the Neubauer Foundation for the PhD fellowship. L.C. and T.A.M.B. thank the support of the HUJI-UK-Spine joint seed funding.

## Authors contributions

J.B., M.G., L.C. and T.A.M.B. designed research. J.B., M.G., C.P. and T.A.M.B. performed research J.B., M.G., C.P., D.A., S.K., L.C. and T.A.M.B. analysed data. J.B., M.G., L.C. and T.A.M.B wrote the manuscript with support from all the authors.

## Methods

### Construction of mutant TasA

For the creation of mutant TasA, corresponding regions of the *tasA* gene were amplified using genomic DNA of the wild-type *B. subtilis* PY79 strain by polymerase chain reaction. Gibson Assembly was utilised to create constructs containing a kanamycin resistance marker and the desired mutation, which were then transformed into PY79. Successful genomic replacements were selected for using kanamycin agar plates, and confirmed using sequencing.

### Transformation protocol

Genomic replacement DNA (gDNA) was transformed into *B. subtilis* strains by picking colonies and growing cultures in 1XMC medium (10 mM Potassium Phosphate buffer pH 7.0, 0.3 mM Sodium Citrate, 0.2% Glucose, 2.2 mg/mL Ferric Ammonium Citrate, 0.01% Casein Hydrolysate, 0.02% Potassium Glutamate) with 0.01 M MgSO_4_ and incubated at 37 °C for 3.5 hours (h) at 250 revolutions per minute (rpm). Then, 300 µl of the culture was mixed with 10 µl of the extracted gDNA and incubated at 37 °C for additional 3 h at 250 rpm. The transformed cells were then plated on selective kanamycin agar plates and incubated at 37 °C for 16 h. Single colonies of each mutant were then picked and used for the TasA purification as described below.

### Native and mutated TasA purification

Native and mutated TasA were purified as described previously [17, 28]. TasA purification was verified using Sodium dodecyl sulphate-polyacrylamide gel electrophoresis (SDS-PAGE) and Western Blot assay (Figures S6B-S6C) and their structure was characterized using circular dichroism (Figure S6G). The concentration of TasA was determined using a BCA protein assay (ThermoFisher Scientific). TasA was passed through a HiLoad 26/60 Superdex S200 sizing column that was pre-equilibrated with 20 mM Tris (pH 8) and lyophilized.

### Circular Dichroism

Circular dichroism (CD) spectra were recorded using a Jasco J-715 spectropolarimeter in the range of 190-260 nm using 1 mm quartz cuvettes. Five scans were recorded and averaged for each sample. CD spectra of buffer solutions were subtracted from the corresponding spectra and ellipticity was converted to mean residue ellipticity (MRE) units using the following equation: 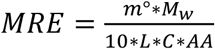, where M_w_ is the molecular weight of the protein in g/mol, L is the cuvette path length in cm, C is the protein concentration in mg/ml and AA in the number of amino acids in the protein sequence. CD was performed using 200 µl of TasA solutions (10 µM of TasA wild-type, 5 µM of Δ28-38 and 174-177_AAAA_ mutants) in 10 mM phosphate buffer, pH 8.0.

### Biofilm formation

Wild-type (NCIB361O), *tasA* mutant ZK3657 (361O tasA::kan) and mutant strains Δ*tasA*, Δ28-38 and *tasA* 174-177_AAAA_ of *B. subtilis* were used for biofilm formation either on agar plates or as pellicles at the liquid-air interface, as specified below. Biofilm formation on agar plates: Biofilms on agar plates were prepared by placing 2 µl of an overnight LB culture on a 1.5% agar MSgg plates [6] and incubating at 30 °C for 3 days. *B. subtilis* Δ*eps* Δ*sinR* biofilms for cryo-EM sample preparation (Figure 5) were incubated at 30 °C for 5 days.

For pellicle formation at the liquid-air interface, pellicles were prepared in 96-well cell culture plates (Costar 3596) by a 1:50 dilution of an overnight LB culture into MSgg broth. Plates were then incubated at 30 °C for 24 h.

### Atomic Force Microscopy

Atomic force microscopy (AFM) topography scans were performed using a BioScope Resolve™ BioAFM (Bruker, Santa Barbara, CA, USA). Height sensor and peak force AFM images were measured in Peak Force Quantitative Nanomechanical Mapping (QNM) mode using ScanAsyst-Air probes (Bruker, resonance frequency 70 KHz, spring constant 0.4 N/m, tip nominal radius 2 nm). Cantilever spring constants were calibrated using the Thermal Tune noise method.

Protein samples were adsorbed on mica as follows: 20 µl of 8 µM TasA in 20 mM Tris pH 8.0, 1.5 M NaCl was placed on a freshly cleaved mica and kept for 30 min in a humid atmosphere. The sample was then washed twice with water for 10 min and dried with a stream of nitrogen.

Pellicle samples were prepared by collecting 24 h-old pellicles (strain ZK5041), resuspending them with 100 µl of 10 mM potassium phosphate buffer pH 8.0, and placing a 20 µl drop on petri dishes (WillCoDish, glass-bottom, Amsterdam) that were pre-incubated with poly-L-lysine. Samples were left to dry on the plate overnight. For poly-L-lysine adsorption on petri dishes, we incubated the petri dishes for 30 minutes with a 0.1 mg/ml poly-L-lysine solution in a humid atmosphere and discarded the poly-L-lysine solution prior to placing a pellicle drop.

### Cryo-EM and cryo-ET sample preparation

For cryo-EM grid preparation of purified TasA fibres, fibre isolates were diluted to final concentrations of 10 µM TasA in 12.5 mM Tris pH 8 and 25 mM NaCl, and 2.5 µl were applied to a freshly glow-discharged Quantifoil R 2/2 Cu/Rh 200 mesh grid and plunge-frozen into liquid ethane using a Vitrobot Mark IV (ThermoFisher) at 100% humidity at 10 °C. For tomography samples of purified TasA, 10 nm Protein-A-gold beads (CMC Utrecht) were added as fiducials prior to plunge-freezing. For cellular cryo-EM sample preparation, *B. subtilis* Δ*eps* Δ*sinR* biofilm was scraped from a plate and resuspended in PBS immediately prior to sample preparation in the same manner.

### Cryo-EM and cryo-ET data collection

Cryo-EM data was collected in a Titan Krios G3 microscope (ThermoFisher) operating at an acceleration voltage of 300 kV, fitted with a Quantum energy filter (slit width 20 eV) and a K3 direct electron detector (Gatan). Images were collected in super-resolution counting mode using a pixel size of 1.092 Å/pixel for helical reconstruction of TasA fibres and 3.402 Å/pixel for imaging of biofilm and mutant TasA samples. For helical reconstruction of TasA, movies were collected as 40 frames, with a total dose of 48.5 electrons/Å^2^, using a range of defoci between -1 and -2 µm. For imaging of *B. subtilis* biofilm ECM components, a total dose of 47.5 e/Å^2^ distributed over 80 frames and defoci between -3 to -7 µm were used. Cryo-ET tilt series of TasA bundles were collected on the same microscope using SerialEM [45], with a total dose of 120-180 e/Å^2^ and defoci of -8 µm, collected between ±60º tilt of the stage at 1º tilt increments.

### Cryo-EM data processing

Helical reconstruction of TasA fibres was performed in RELION [46, 47]. Movies were corrected using the RELION 3.1 implementation of MotionCor2 [48], and CTF parameters were estimated using CTFFIND4 [49]. Three-dimensional classification was used to identify a subset of particles that supported refinement to 3.5 Å resolution. Symmetry searches were used during reconstruction, resulting in a final rise of 48.36 Å and a left-handed twist per subunit of 55.2°. Resolution was estimated using the gold-standard FSC method as implemented in RELION. Local resolution measurements were also performed using RELION.

### Model building and refinement

Manual model building was performed in Coot [50]. The initial model of fibrous TasA was subjected to real-space refinement against the cryo-EM map within the Phenix package [51]. Three subunits of TasA were built and used for final refinement. Non-crystallographic symmetry between individual TasA subunits was applied for all refinement runs. Model validation including map-vs-model resolution estimation was performed in Phenix.

### Tomogram reconstruction and fitting of atomic models

Tilt series alignment via tracking of gold fiducials was performed using the etomo package as implemented in IMOD [52]. Tomograms were reconstructed with WBP in IMOD [52] or SIRT in tomo3d [53]. Atomic models of TasA fibres containing eighteen subunits were fitted in parallel into the tomographic density using ChimeraX [54]. Fits consistently placed a loop containing residues 174-177, which faces outwards from the helical centre, at the inter-subunit interface, independent of fibre orientation.

### Molecular Modelling and Molecular Dynamics Simulations

The cryo-EM structure of a TasA filament doublet was submitted to the Martini Maker tool [55] from the CHARMM-GUI server [56, 57] to build the initial coarse-grained model of the system. System was solvated with explicit water, and a number of counter ions were added to neutralize charges, with an extra salt concentration of 1 M of KCl ions for all simulations. The final system obtained was composed of approximately 2.5 million particles. Molecular dynamics simulations were performed using the GROMACS simulation suite (version 2020.4) [58] along with the Martini2.2 force field [59]. The ElNeDyn model with an elastic network cut-off of 0.9 nm and a force constant of 500 kJ.mol^−1^.nm^−2^ was used to coarse-grain the protein models [60]. Starting positions for the filaments were oriented along the Y axis and over the periodic boxes in a way that allowed the extremities of each filament to interact non-covalently with their periodic images, forming a continuous and stable structure.

Two stages of equilibration were performed employing the NVT and NPT ensembles each for 10 ns while keeping the protein beads restrained but allowing the water and ions to relax. Two independent production simulations each of duration 200 ns with were then performed at 313 K using the velocity rescale thermostat [61] with a coupling constant of τ = 1 Pressure coupling was implemented with the Parrinello-Rahman barostat [61] with a time constant of 12 ps. Electrostatics were treated with the reaction-field method, with dielectric constants of 15 and infinity for charge screening in both short- and long-range regimes, respectively. The cut-off for non-bonded and electrostatic interactions in the short-range was 1.1 nm. Starting random velocities were modified at the beginning of each replicate to improve conformational sampling. Molecules were manipulated, visualized, and analysed utilizing VMD [62] and Pymol [63] software. Quantification of the interaction between filaments was performed by calculating the time any residue from any of the filaments was within a distance of 0.6 nm from each other. The interaction time of each binding-pair of residues was converted to a scale of 0 to 1, where 0 = 0% and 1 = 100% of time spent within the cut-off.

### Alphafold 2 prediction of a TapA/TasA complex

The TapA/TasA complex was predicted using AlphaFold 2.1 [34]. Signal peptide cleavage for TapA was predicted with SignalP [64]. Sequence alignment of the mature forms of TapA and TasA was performed using the Clustal Omega service provided by EMBL-EBI [65].

### Data visualisation and quantification

Cryo-EM images were visualised in IMOD. Fiji [66] was used for bandpass and Gaussian filtering, followed by automatic contrast adjustment. Atomic structures and tomographic data were displayed in ChimeraX. To analyse the effect of amino acid residues 28-38 truncation on fibre formation, a random subset of 20 micrographs was chosen using the *shuf* functionality of GNU, and TasA fibres were counted manually in each micrograph. For quantifying bundling in the TasA 174-177_AAAA_ mutant, fibres with the same periodicity as the unaltered TasA fibres were counted and considered bundling if there were at least two visible periodic fibre interactions (see Figures 4B-C); 100 fibres were counted from randomly selected micrographs. For homology modelling of CalY1 (Figure S3), SWISS-MODEL [67] server was used with the cryo-EM structure of the TasA fibre as the template model.

## Supplementary Figures

**Figure S1.**
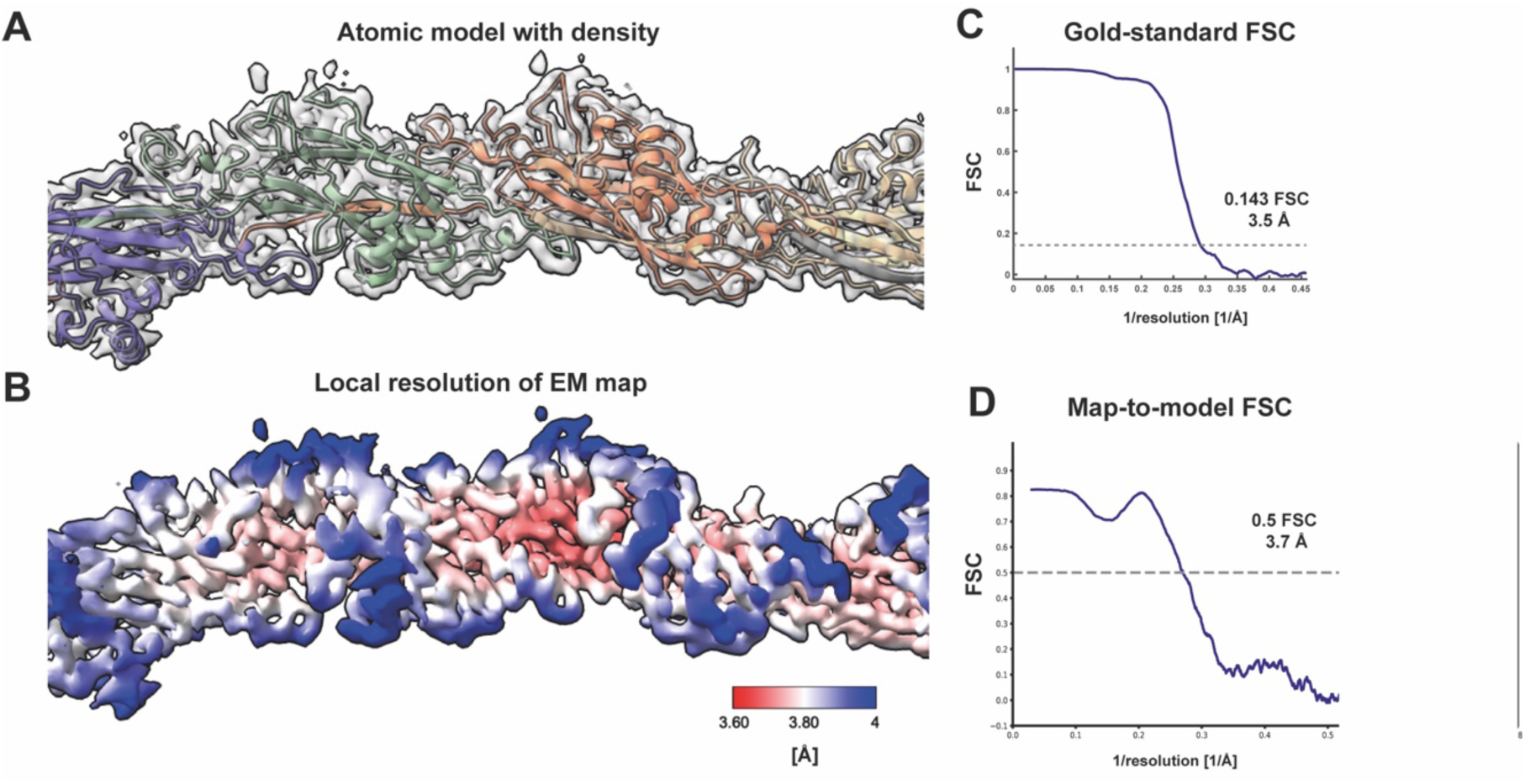
Cryo-EM structure of TasA fibres. **(A)** TasA fibre atomic model, shown as differently coloured ribbon diagrams, within the cryo-EM density shown at 12 σ isosurface contour level. **(B)** Local resolution estimation of the cryo-EM reconstruction. **(C)** Gold-standard Fourier Shell Correlation (FSC) plot. **(D)** Map-to-model FSC curve.

**Figure S2.**
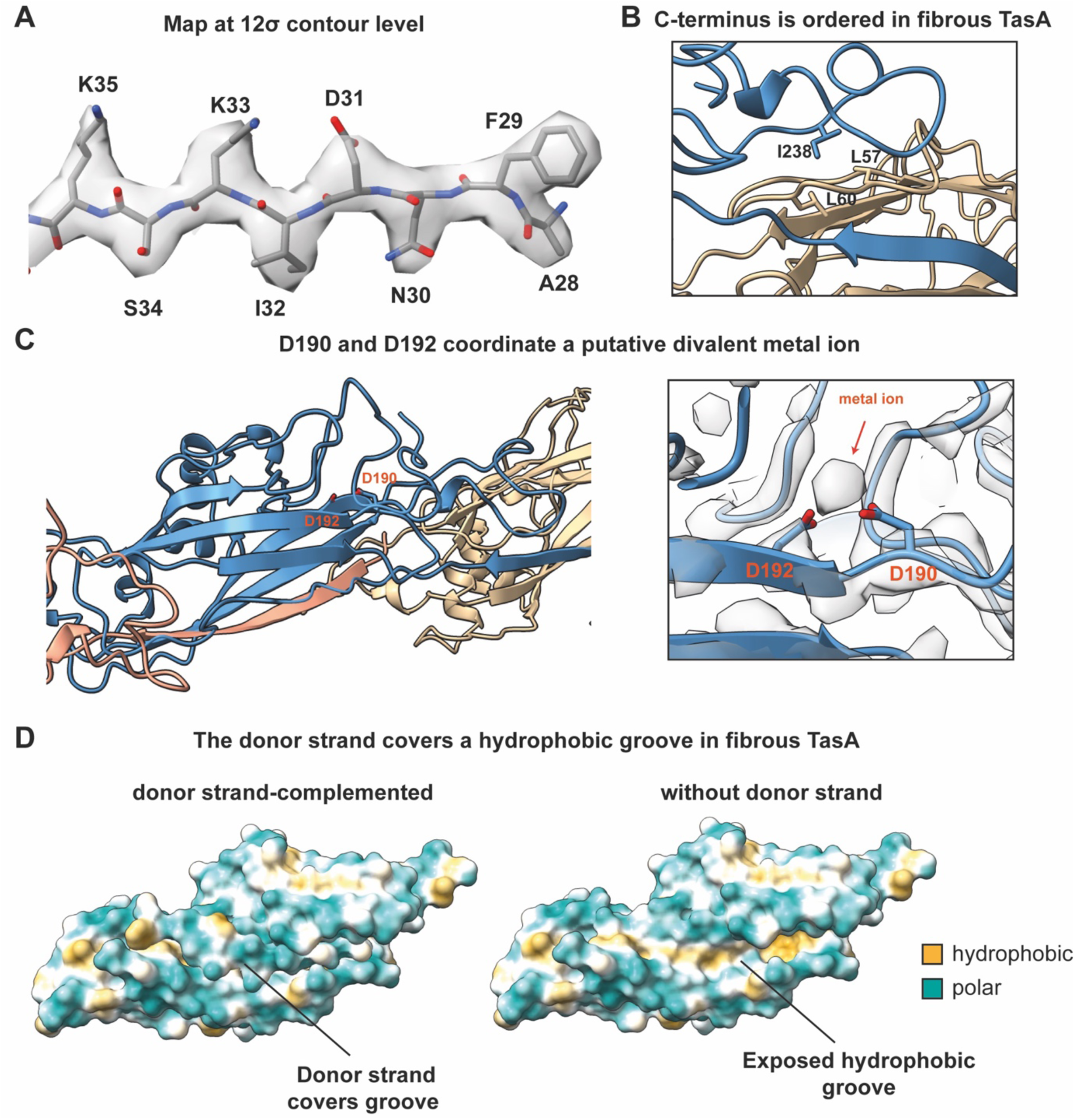
Structural features of TasA. **(A)** Map at a 12σ contour level with atomic model of TasA N-terminal residues shown. **(B)** Interaction of hydrophobic residue I238 of the n_th_ subunit (blue ribbon) within the C-terminus, with the (n+1)_th_ subunit (beige ribbon). **(C)** A putative metal ion co-ordination site is resolved in the cryo-EM map. While cryo-EM does not allow determining the chemical identity of the ion, TasA shows homology to the camelysin CalY in *B. cereus*, which was found to contain a zinc ion [68]. **(D)** The donor strand shields a hydrophobic groove in TasA (left), which is exposed in the absence of the donor strand (right).

**Figure S3.**
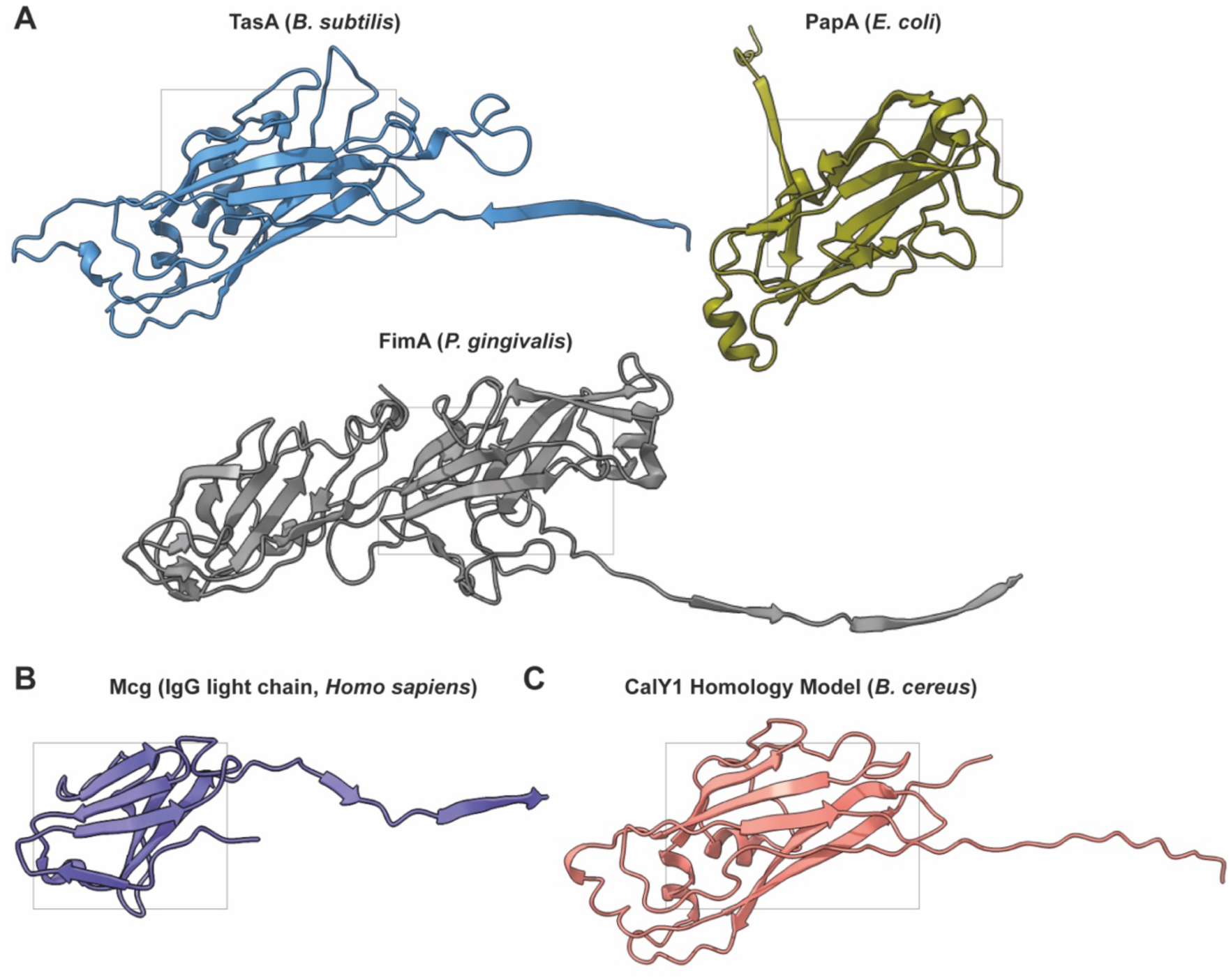
Comparison of the TasA structure (fibre form) with proteins from other systems undergoing donor-strand complementation. **(A)** Comparison of TasA (blue ribbon) with the PapA Chaperone-Usher Pathway pilin of a type I pilus from *Escherichia coli* [69] (PDB 5FLU, military green ribbon) and the FimA pilin of a type V pilus from *Porphyromonas gingivalis* [32] (PDB 6KMF, grey ribbon). **(B)** Donor strand-complemented structure of the Mcg IgG light chain protein involved in amyloidosis. **(C)** Based on the atomic structure of TasA, we also created a homology model of an accessory protein called CalY1 which is encoded in the same operon as TasA in *B. cereus* [68, 70]. Given significant sequence homology to TasA and conservation of hydrophobic residues involved in donor strand complementation, CalY1 might also polymerise using a donor strand mechanism. These comparisons suggest that a β-sandwich (grey boxes) fold is a common structural feature of proteins undergoing donor strand complementation.

**Figure S4.**
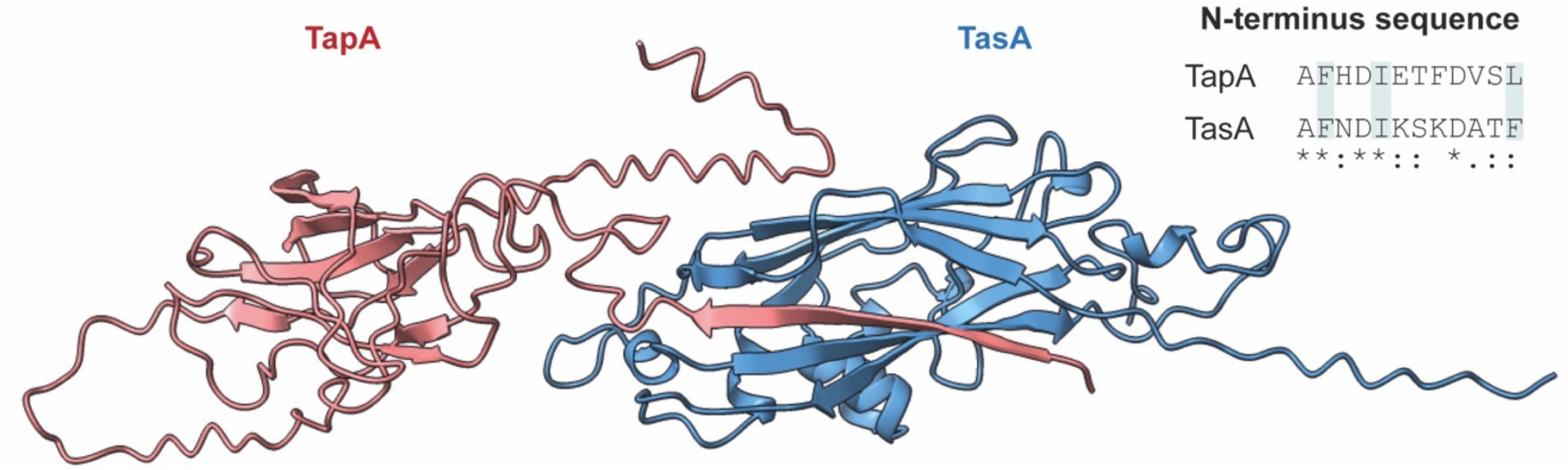
TapA is predicted to extend a donor strand that complements TasA. Shown is an AlphaFold 2 multimer prediction of a TapA/TasA complex. A Clustal Omega alignment of N-terminal residues of TapA and TasA is shown on the upper right; the hydrophobic residues mediating major donor strand interactions with the complemented subunit (see Figure 1) are marked in cyan. Asterisks indicate full conservation, colons indicate conservation between groups of strongly similar properties, and periods indicate conservation between groups of weakly similar properties as per Clustal Omega output.

**Figure S5.**
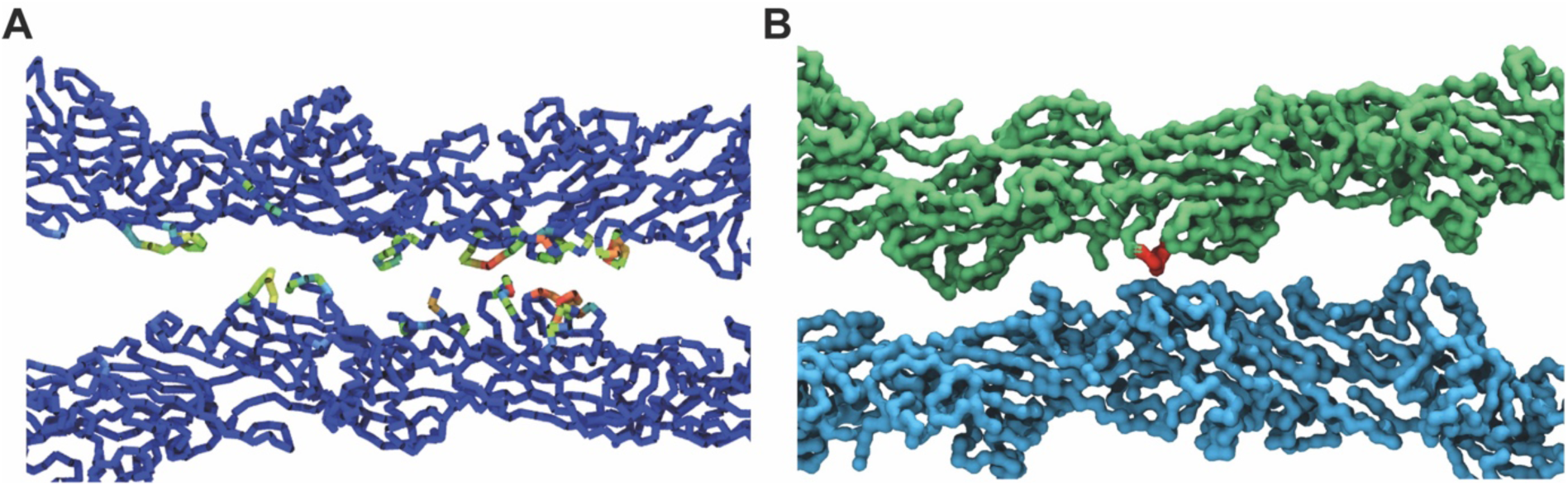
MD simulations of TasA doublets. **(A)** Stable solution of coarse-grained MD (200 ns) of two interacting TasA fibres with residues found interacting during the simulation at the inter-fibre interface; red: strong interactions, green, weak interactions. Residues found to be interacting are 153-160 (IKKQIDPK), 173-179 (IDKGTAP), 185-190 (PKTPTD), 238-240 (IKK), 243-245 (TDK), and 248 (Y). **(B)** Residues 174-177 are marked red for comparison in the same doublet model from MD.

**Figure S6.**
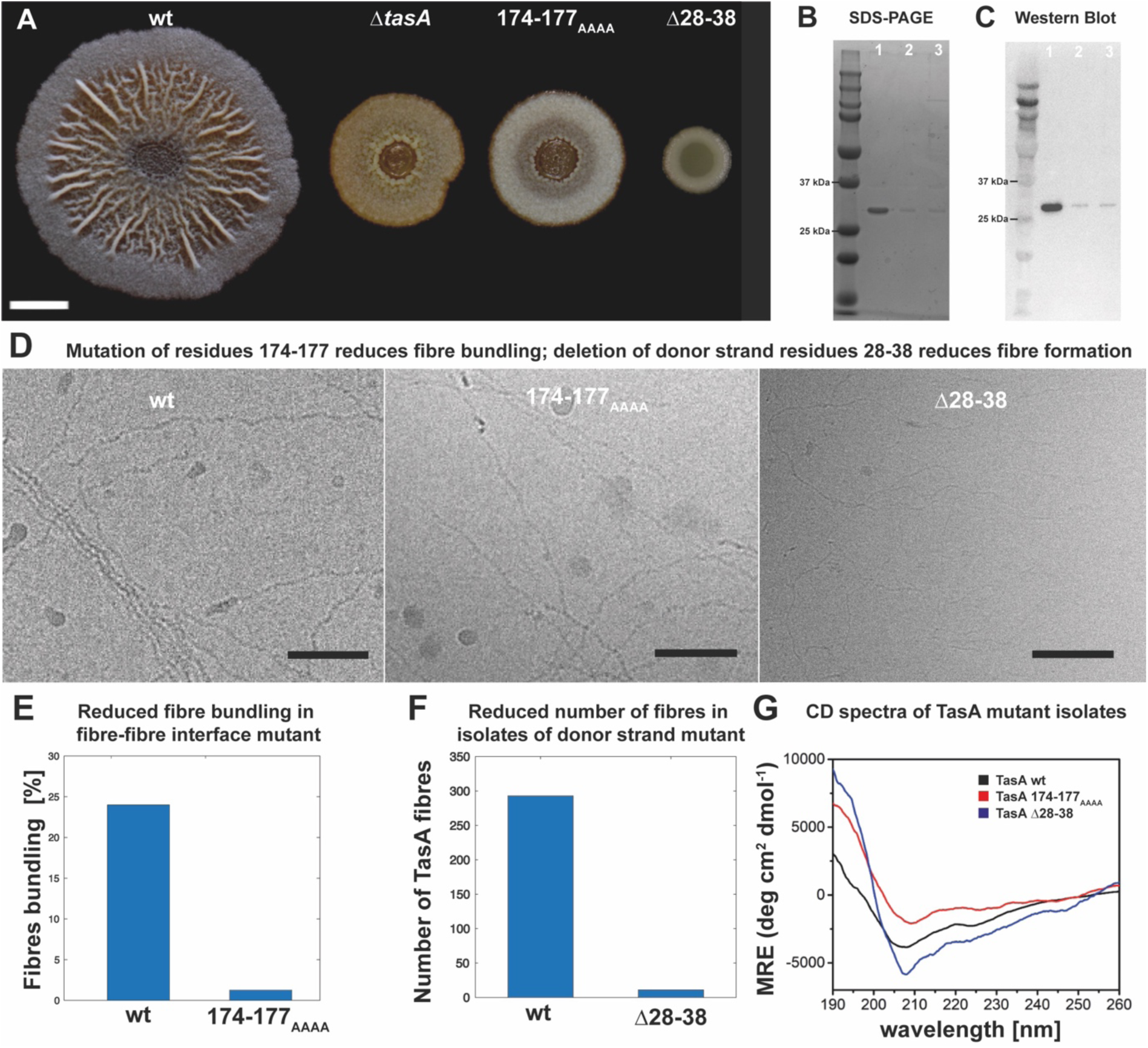
Mutational studies of TasA. **(A)** *B. subtilis* biofilms grown on agar plates, imaged after 72 hours post inoculation. From left to right: wild-type (wt), Δ*tasA, tasA* 174-177_AAAA_ (fibre-fibre interface mutant) and *tasA* Δ28-38 (donor strand mutant). Scale bar corresponds to 0.5 cm. **(B-C)** SDS-PAGE and Western blot of TasA protein isolates. Lane 1: wt TasA, lane 2: TasA Δ28-38, lane 3: TasA 174-177_AAAA_, **(D)** cryo-EM images comparing wild-type with mutant TasA. In a fibre-fibre interface mutant TasA (174-177_AAAA_ TasA, middle), bundling is significantly reduced. A donor strand mutant TasA (Δ28-38 TasA, right) shows a markedly reduced number of TasA fibres, while smaller aberrant, contaminating fibres are visible instead. All specimens are at the same measured protein concentrations. **(E)** Quantification of bundling behaviour in the fibre-fibre interface mutant TasA (174-177_AAAA_). **(F)** Quantification of TasA fibres in cryo-EM images of donor strand mutant TasA (Δ28-38). **(G)** Circular Dichroism spectra of unmutated, wt TasA (purified from Δ*sinR* Δ*eps* strain), 174-177_AAAA_ TasA and Δ28-38 TasA.

**Figure S7.**
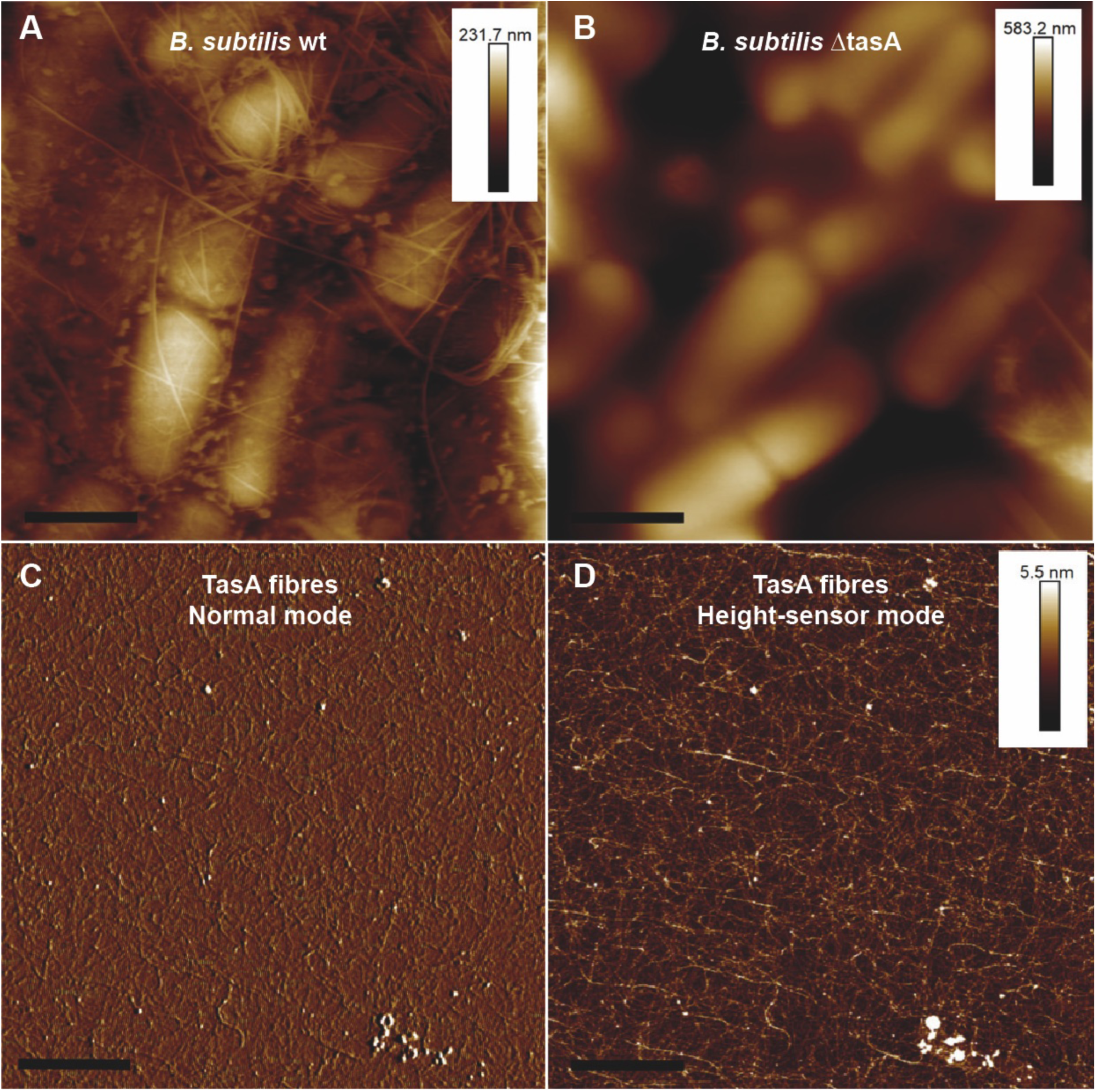
AFM imaging of *B. subtilis* biofilms and purified TasA fibres. **(A-B)** Height sensor mode AFM images of (A) wt and (B) Δ*tasA* pellicles. **(C)** AFM peak force error and **(D)** height sensor mode AFM images of TasA fibres formed *in vitro* from purified protein. Height scale is shown to the top-right of each relevant image. Scale bars corresponds to 1 µm.

**Figure S8.**
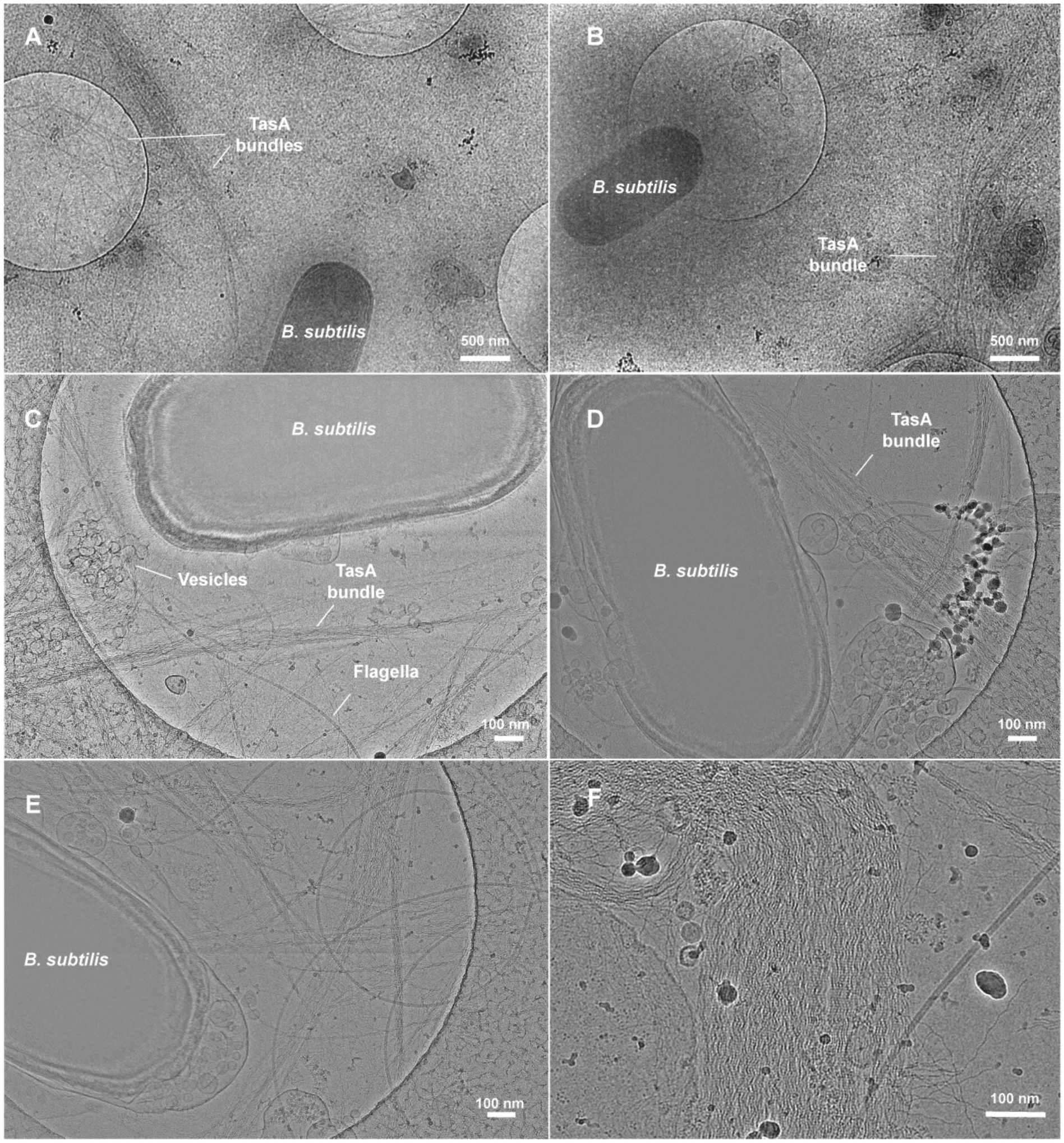
Cryo-EM of the ECM components of *B. subtilis* Δ*eps* Δ*sinR* biofilms. **(A-F)** Gallery of medium- and high magnification images of resuspended *B. subtilis* Δ*eps* Δ*sinR* biofilms shows TasA bundles surrounding *B. subtilis* cells.

## References

1. O’Toole, G., H.B. Kaplan, and R. Kolter, Biofilm formation as microbial development. Annual Reviews in Microbiology, 2000. 54(1): p. 49–79.

2. Donlan, R.M., Biofilms: microbial life on surfaces. Emerging Infectious Diseases, 2002. 8(9): p. 881.

3. Flemming, H.-C., et al., Biofilms: an emergent form of bacterial life. Nature Reviews Microbiology, 2016. 14(9): p. 563–575.

4. Mah, T.-F.C. and G.A. O’Toole, Mechanisms of biofilm resistance to antimicrobial agents. Trends in Microbiology, 2001. 9(1): p. 34–39.

5. Flemming, H.-C. and J. Wingender, The biofilm matrix. Nature Reviews Microbiology, 2010. 8(9): p. 623–633.

6. Branda, S.S., et al., Fruiting body formation by Bacillus subtilis. Proceedings of the National Academy of Sciences, 2001. 98(20): p. 11621–11626.

7. Chen, Y., et al., A Bacillus subtilis sensor kinase involved in triggering biofilm formation on the roots of tomato plants. Molecular Microbiology, 2012. 85(3): p. 418–430.

8. Chu, F., et al., A novel regulatory protein governing biofilm formation in Bacillus subtilis. Molecular Microbiology, 2008. 68(5): p. 1117–1127.

9. Chai, Y., et al., Galactose metabolism plays a crucial role in biofilm formation by Bacillus subtilis. MBio, 2012. 3(4): p. e00184–12.

10. Kobayashi, K., SlrR/SlrA controls the initiation of biofilm formation in Bacillus subtilis. Molecular Microbiology, 2008. 69(6): p. 1399–1410.

11. Azulay, D.N., et al., Multiscale X-ray study of Bacillus subtilis biofilms reveals interlinked structural hierarchy and elemental heterogeneity. Proceedings of the National Academy of Sciences, 2022. 119(4): p. e2118107119.

12. Vlamakis, H., et al., Sticking together: building a biofilm the Bacillus subtilis way. Nature Reviews Microbiology, 2013. 11(3): p. 157–168.

13. Branda, S.S., et al., A major protein component of the Bacillus subtilis biofilm matrix. Molecular Microbiology, 2006. 59(4): p. 1229–1238.

14. Stöver, A.G. and A. Driks, Secretion, localization, and antibacterial activity of TasA, a Bacillus subtilis spore-associated protein. Journal of Bacteriology, 1999. 181(5): p. 1664–1672.

15. Chu, F., et al., Targets of the master regulator of biofilm formation in Bacillus subtilis. Molecular Microbiology, 2006. 59(4): p. 1216–1228.

16. Kearns, D.B., et al., A master regulator for biofilm formation by Bacillus subtilis. Molecular Microbiology, 2005. 55(3): p. 739–749.

17. Romero, D., et al., Amyloid fibers provide structural integrity to Bacillus subtilis biofilms. Proceedings of the National Academy of Sciences, 2010. 107(5): p. 2230–2234.

18. Romero, D., et al., Functional analysis of the accessory protein TapA in Bacillus subtilis amyloid fiber assembly. Journal of Bacteriology, 2014. 196(8): p. 1505–1513.

19. Romero, D., et al., An accessory protein required for anchoring and assembly of amyloid fibres in B. subtilis biofilms. Molecular Microbiology, 2011. 80(5): p. 1155–1168.

20. Terra, R., et al., Identification of Bacillus subtilis SipW as a bifunctional signal peptidase that controls surface-adhered biofilm formation. Journal of Bacteriology, 2012. 194(11): p. 2781–2790.

21. Diehl, A., et al., Structural changes of TasA in biofilm formation of Bacillus subtilis. Proceedings of the National Academy of Sciences, 2018. 115(13): p. 3237–3242.

22. Cámara-Almirón, J., et al., Dual functionality of the amyloid protein TasA in Bacillus physiology and fitness on the phylloplane. Nature Communications, 2020. 11(1): p. 1–21.

23. Mammeri, N.E., et al., Molecular architecture of bacterial amyloids in Bacillus biofilms. The FASEB Journal, 2019. 33(11): p. 12146–12163.

24. Balistreri, A., E. Goetzler, and M. Chapman, Functional amyloids are the rule rather than the exception in cellular biology. Microorganisms, 2020. 8(12): p. 1951.

25. Chapman, M.R., et al., Role of Escherichia coli curli operons in directing amyloid fiber formation. Science, 2002. 295(5556): p. 851–855.

26. Dueholm, M.S., et al., Functional amyloid in Pseudomonas. Molecular Microbiology, 2010. 77(4): p. 1009–1020.

27. Erskine, E., et al., Formation of functional, non-amyloidogenic fibres by recombinant Bacillus subtilis TasA. Molecular Microbiology, 2018. 110(6): p. 897–913.

28. Chai, L., et al., Isolation, characterization, and aggregation of a structured bacterial matrix precursor. Journal of Biological Chemistry, 2013. 288(24): p. 17559–17568.

29. Ghrayeb, M., et al., Fibrilar Polymorphism of the Bacterial Extracellular Matrix Protein TasA. Microorganisms, 2021. 9(3): p. 529.

30. Greenwald, J. and R. Riek, Biology of amyloid: structure, function, and regulation. Structure, 2010. 18(10): p. 1244–1260.

31. Remaut, H., et al., Donor-strand exchange in chaperone-assisted pilus assembly proceeds through a concerted β strand displacement mechanism. Molecular Cell, 2006. 22(6): p. 831–842.

32. Shibata, S., et al., Structure of polymerized type V pilin reveals assembly mechanism involving protease-mediated strand exchange. Nature Microbiology, 2020. 5(6): p. 830–837.

33. Brumshtein, B., et al., Formation of amyloid fibers by monomeric light chain variable domains. Journal of Biological Chemistry, 2014. 289(40): p. 27513–27525.

34. Jumper, J., et al., Highly accurate protein structure prediction with AlphaFold. Nature, 2021. 596(7873): p. 583–589.

35. Earl, C., et al., The majority of the matrix protein TapA is dispensable for Bacillus subtilis colony biofilm architecture. Molecular Microbiology, 2020. 114(6): p. 920–933.

36. Bharat, T.A., et al., Structures of actin-like ParM filaments show architecture of plasmid-segregating spindles. Nature, 2015. 523(7558): p. 106–110.

37. Sleutel, M., B. Pradhan, and H. Remaut, Structural analysis of the bacterial amyloid curli. bioRxiv, 2022: p. 2022.02.28.482343.

38. Cucarella, C., et al., Bap, a Staphylococcus aureus surface protein involved in biofilm formation. Journal of Bacteriology, 2001. 183(9): p. 2888–2896.

39. Tayeb-Fligelman, E., et al., The cytotoxic Staphylococcus aureus PSMα3 reveals a cross-α amyloid-like fibril. Science, 2017. 355(6327): p. 831–833.

40. Berk, V., et al., Molecular architecture and assembly principles of Vibrio cholerae biofilms. Science, 2012. 337(6091): p. 236–9.

41. Melia, C.E., et al., Architecture of cell-cell junctions in situ reveals a mechanism for bacterial biofilm inhibition. Proc Natl Acad Sci U S A, 2021. 118(31).

42. Buzzo, J.R., et al., Z-form extracellular DNA is a structural component of the bacterial biofilm matrix. Cell, 2021. 184(23): p. 5740-5758. e17.

43. Tarafder, A.K., et al., Phage liquid crystalline droplets form occlusive sheaths that encapsulate and protect infectious rod-shaped bacteria. Proceedings of the National Academy of Sciences, 2020. 117(9): p. 4724–4731.

44. Secor, P.R., et al., Filamentous bacteriophage promote biofilm assembly and function. Cell Host & Microbe, 2015. 18(5): p. 549–559.

45. Mastronarde, D.N., Automated electron microscope tomography using robust prediction of specimen movements. Journal of Structural Biology, 2005. 152(1): p. 36–51.

46. Scheres, S.H., RELION: implementation of a Bayesian approach to cryo-EM structure determination. Journal of Structural Biology, 2012. 180(3): p. 519–530.

47. He, S. and S.H. Scheres, Helical reconstruction in RELION. Journal of Structural Biology, 2017. 198(3): p. 163–176.

48. Zheng, S.Q., et al., MotionCor2: anisotropic correction of beam-induced motion for improved cryo-electron microscopy. Nature Methods, 2017. 14(4): p. 331–332.

49. Rohou, A. and N. Grigorieff, CTFFIND4: Fast and accurate defocus estimation from electron micrographs. Journal of Structural Biology, 2015. 192(2): p. 216–221.

50. Emsley, P. and K. Cowtan, Coot: model-building tools for molecular graphics. Acta Crystallographica Section D: Biological Crystallography, 2004. 60(12): p. 2126–2132.

51. Adams, P.D., et al., PHENIX: a comprehensive Python-based system for macromolecular structure solution. Acta Crystallographica Section D: Biological Crystallography, 2010. 66(2): p. 213–221.

52. Kremer, J.R., D.N. Mastronarde, and J.R. McIntosh, Computer visualization of three-dimensional image data using IMOD. Journal of Structural Biology, 1996. 116(1): p. 71–76.

53. Agulleiro, J.-I. and J.-J. Fernandez, Tomo3D 2.0–exploitation of advanced vector extensions (AVX) for 3D reconstruction. Journal of Structural Biology, 2015. 189(2): p. 147–152.

54. Goddard, T.D., et al., UCSF ChimeraX: Meeting modern challenges in visualization and analysis. Protein Science, 2018. 27(1): p. 14–25.

55. Hsu, P.C., et al., CHARMM-GUI Martini Maker for modeling and simulation of complex bacterial membranes with lipopolysaccharides. Journal of Computational Chemistry, 2017.

56. Jo, S., et al., CHARMM-GUI: a web-based graphical user interface for CHARMM. Journal of Computational Chemistry, 2008. 29(11): p. 1859–1865.

57. Lee, J., et al., CHARMM-GUI input generator for NAMD, GROMACS, AMBER, OpenMM, and CHARMM/OpenMM simulations using the CHARMM36 additive force field. Journal of Chemical Theory and Computation, 2016. 12(1): p. 405–413.

58. Abraham, M.J., et al., GROMACS: High performance molecular simulations through multi-level parallelism from laptops to supercomputers. SoftwareX, 2015. 1: p. 19–25.

59. de Jong, D.H., et al., Improved parameters for the martini coarse-grained protein force field. Journal of Chemical Theory and Computation, 2013. 9(1): p. 687–697.

60. Periole, X., et al., Combining an elastic network with a coarse-grained molecular force field: structure, dynamics, and intermolecular recognition. Journal of Chemical Theory and Computation, 2009. 5(9): p. 2531–2543.

61. Bussi, G., D. Donadio, and M. Parrinello, Canonical sampling through velocity rescaling. The Journal of Chemical Physics, 2007. 126(1): p. 014101.

62. Humphrey, W., A. Dalke, and K. Schulten, VMD: visual molecular dynamics. Journal of Molecular Graphics, 1996. 14(1): p. 33–38.

63. Schrodinger, L., The PyMOL molecular graphics system. Version, 2010. 1(5): p. 0.

64. Armenteros, J.J.A., et al., SignalP 5.0 improves signal peptide predictions using deep neural networks. Nature Biotechnology, 2019. 37(4): p. 420–423.

65. Madeira, F., et al., The EMBL-EBI search and sequence analysis tools APIs in 2019. Nucleic Acids Research, 2019. 47(W1): p. W636–W641.

66. Schindelin, J., et al., Fiji: an open-source platform for biological-image analysis. Nature Methods, 2012. 9(7): p. 676–682.

67. Schwede, T., et al., SWISS-MODEL: an automated protein homology-modeling server. Nucleic Acids Research, 2003. 31(13): p. 3381–3385.

68. Grass, G., et al., Camelysin is a novel surface metalloproteinase from Bacillus cereus. Infection and Immunity, 2004. 72(1): p. 219–228.

69. Hospenthal, M.K., et al., Structure of a chaperone-usher pilus reveals the molecular basis of rod uncoiling. Cell, 2016. 164(1-2): p. 269–278.

70. Caro-Astorga, J., et al., A genomic region involved in the formation of adhesin fibers in Bacillus cereus biofilms. Frontiers in Microbiology, 2015. 5: p. 745.

